# A novel Bayesian method for inferring and interpreting the dynamics of adaptive landscapes from phylogenetic comparative data

**DOI:** 10.1101/004465

**Authors:** Josef C. Uyeda, Luke J. Harmon

## Abstract

Our understanding of macroevolutionary patterns of adaptive evolution has greatly increased with the advent of large-scale phylogenetic comparative methods. Widely used Ornstein-Uhlenbeck (OU) models can describe an adaptive process of divergence and selection. However, inference of the dynamics of adaptive landscapes from comparative data is complicated by interpretational difficulties, lack of identifiability among parameter values and the common requirement that adaptive hypotheses must be assigned *a priori*. Here we develop a reversible-jump Bayesian method of fitting multi-optima OU models to phylogenetic comparative data that estimates the placement and magnitude of adaptive shifts directly from the data. We show how biologically informed hypotheses can be tested against this inferred posterior of shift locations using Bayes Factors to establish whether our *a priori* models adequately describe the dynamics of adaptive peak shifts. Furthermore, we show how the inclusion of informative priors can be used to restrict models to biologically realistic parameter space and test particular biological interpretations of evolutionary models. We argue that Bayesian model-fitting of OU models to comparative data provides a framework for integrating of multiple sources of biological data—such as microevolutionary estimates of selection parameters and paleontological timeseries—allowing inference of adaptive landscape dynamics with explicit, process-based biological interpretations.

## Introduction

The phenotypic adaptive landscape has been widely used as the conceptual foundation for studying phenotypic evolution across micro- to macroevolutionary scales(Arnold et al. 2001). The concept has been applied to microevolutionary studies of selection (Lande and Arnold 1983), studies of paleontological time-series (Simpson 1944, 1953; Hunt et al. 2008; Reitan et al. 2012) and to stochastic models of trait evolution fit to phylogenetic comparative data (Hansen 1997; Butler and King 2004; Hansen et al. 2008; Uyeda et al. 2011; Eastman et al. 2013). Consequently, the adaptive landscape has the potential to unite micro to macroevolution into a single cohesive theoretical framework (Arnold et al. 2001; Hansen 2012). However, a major disconnect between microevolutionary and macroevolutionary formulations of adaptive landscapes is that microevolutionary studies typically examine static landscapes, whereas macroevolutionary patterns result from the dynamics of adaptive peak movement over long evolutionary timescales (Gavrilets 2004; Hansen 2012). While macroevolutionary models fit to phylogenetic comparative data almost certainly describe the cumulative dynamics of adaptive landscapes, these phenomenological models are disconnected from adaptive landscapes at shorter timescales, and thus, become difficult to interpret in terms of biological processes. Synthesis will require a unification of theory and data across scales that allows inference of the dynamics of the movement of adaptive landscapes directly from macroevolutionary data (Uyeda et al. 2011).

Existing models of adaptive evolution at macroevolutionary scales typically rely the Ornstein-Uhlenbeck (OU) model of trait evolution (Hansen 1997; Butler and King 2004), which has a strong connection to the concept of adaptive landscapes (Lande 1976). Fitting OU models to macroevolutionary data allows researchers to test hypotheses regarding the existence of distinct phenotypic optima between groups of species (Butler and King 2004; Beaulieu et al. 2012) and more generally, infer evolutionary regressions between phenotypic traits and predictor variables (Hansen et al. 2008; Hansen and Bartoszek 2012). Hansen (1997) introduced the method to phylogenetic comparative methods as a means to test specific adaptive hypotheses–such as the hypothesis that phenotypic optima for browsing vs. grazing horses are different. Butler and King (2004) extended this method to test among competing adaptive hypotheses and provided a widely used implementation as an R package (

~~~
ouch
~~~

). However, 

~~~
ouch
~~~

 and other software typically require *a priori* assignment of adaptive hypotheses to the phylogeny, with optima “painted” onto branches according to the researcher’s pre-existing hypotheses (but see Ingram and Mahler 2013). Following model-fitting by maximum likelihood, model selection is used among hypothesized scenarios and the best-fitting model is chosen (Butler and King 2004). However, it can be difficult to ascertain whether the specific hypothesis chosen by the researcher is a good hypothesis, or simply better than the tested alternatives. Furthermore, to infer the dynamics of adaptive landscapes themselves, we do not wish to assume a limited set of hypotheses *a priori*, but to estimate the dynamics of phenotypic optima directly from the data. This of course does not mean that we want to throw away biologically informed hypotheses. Rather, we seek a framework for evaluating whether *a priori* hypotheses adequately describe the statistical patterns in the data.

Incorporating biological realism into comparative models remains a challenging goal. Statistical models fit to phylogenies are inherently phenomenological, and may be consistent with multiple biological interpretations. For example, Brownian motion (BM) processes can result from neutral genetic drift of trait means, neutral drift of adaptive peaks, or drifting adaptive zones (Lande 1976; Felsenstein 1985, 1988). Rate tests derived from these models have generally rejected genetic drift as a model for macroevolutionary patterns (Turelli et al. 1988; Lynch 1990; Hohenlohe and Arnold 2008), and simple drift of adaptive peaks seems inconsistent with observed microevolutionary and macroevolutionary data (Estes and Arnold 2007; Uyeda et al. 2011). As with BM models, OU models have an explicitly microevolutionary interpretation in terms of the process of stabilizing selection and genetic drift on static adaptive landscapes (Lande 1976). Alternatively, these optima may represent adaptive zones within which adaptive peaks drift stochastically, or even broad ranges around which dynamically evolving adaptive zones evolve (Hansen 1997). The boundaries between these latter interpretations can become fuzzy, and while a tendency for a species to return to an optimal state is suggestive of adaptive evolution, it remains unclear how fitted models reflect evolutionary processes and patterns. Because of this difficulty, previous authors have tended to use terms such as “adaptive regimes” or “selective regimes” to describe optima fit via OU modeling. These terms only vaguely connect model parameters to the process of adaptive evolution. More specific interpretations of these models will require methods that can more explicitly connect to microevolutionary and paleontological data.

We present a Bayesian framework for studying adaptive evolution using multi-optima OU models that attempts to provide solutions to these challenges. We implement a reversible-jump algorithm that jointly estimates the location, number and magnitude of shifts in adaptive optima from phylogenetic comparative data (Green 1995; Huelsenbeck et al. 2000, 2004; Eastman et al. 2011, 2013; Rabosky et al. 2013). We use simulations to demonstrate the effectiveness of this method at identifying the location of shifts and estimating parameters. Furthermore, we demonstrate how the method can be used to compare specific hypotheses, and how informative priors can be incorporated from different data sources to test mechanistic interpretations of phylogenetic patterns of trait divergence. We incorporate these methods into a flexible software package, 

~~~
bayou
~~~

, for the R statistical environment (R Core Team 2014).

Our approach has a number of distinct advantages over existing methods. First, the reversible-jump framework will produce a full posterior of credible models and parameter values and therefore incorporates uncertainty in regime number, placement and parameter estimates. This is particularly important when modeling OU processes, as these models often have flat ridges on likelihood space, particularly for the correlated parameters *α* and *σ*^2^ (Ho and Ané 2013). However, some parameter values along these ridges are inconsistent with particular biological interpretations of the model. A natural way of restricting model exploration to interesting regions of parameter space is to place priors on the parameters. We compare model fits between biologically informed *a priori* hypotheses of adaptive evolution against the full posterior of credible models using marginal likelihoods. From these comparisons, we can conclude whether a particular hypothesis captures the relevant signal of adaptive peak movement in the data. Furthermore, we show how alternative priors and parameterizations of OU models based on different biological interpretations can be compared, and suggest how additional sources of data may be incorporated into analyses. By using informative priors, we can incorporate data on microevolutionary biological processes (e.g. strength of natural selection, population size, genetic variance) and/or constrain the dynamics of the model to be consistent with patterns observed at other biological scales (e.g. stasis, rapid evolution over short timescales, etc.). By doing so, we obtain a clearer picture of what our comparative models are actually measuring and how we can interpret macroevolutionary patterns.

## Methods

We model phenotypic evolution across phylogenies as an Ornstein-Uhlenbeck (OU) process, which is a mean-reverting continuous-time stochastic process with three parameters describing per unit time magnitude of uncorrelated diffusion (*σ*^2^), the rate of adaptation (*α*) and the optimum value of the process (*θ*) according to the stochastic differential equation:

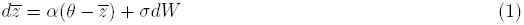

where dW is a continuous-time Wiener process and 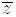 is the trait mean. If *α* = 0, the process reduces to a Brownian motion model of trait evolution. The parameter *α* is measured in inverse time units while the parameter *σ*^2^ is in units of squared trait units per unit time. An easier way to interpret *α* is by reparameterizing the value as the phylogenetic half-life (Hansen et al. 2008), which is defined as *ln*(2)/*α* and is measured in time units. It can be interpreted as the amount of time it takes for the expected trait value to get halfway to the phenotypic optimum. Small values of phylogenetic half-life (i.e. large values of *α*) also have the effect of eroding covariance between species following speciation. A model with a phylogenetic half-life much greater than tree height will thus resemble a BM process, whereas a phylogenetic half-life much shorter than the youngest split on a phylogeny will resemble a white noise process (i.e. residual trait values are completely uncorrelated). We will refer to both *α* and phylogenetic half-life throughout the manuscript, as doing so will make certain patterns more clear and interpretable. In addition, a useful compound parameter for OU processes is the stationary variance [*V*_*y*_ = *σ*^2^/(2*α*)], which is the equilibrium variance of an OU process evolving around a stationary optimum *θ*.

Multi-optima OU models are modifications of the standard OU model in which adaptive optimum, *θ*, varies across the phylogeny according to discrete shifts in adaptive regimes (Hansen 1997; Butler and King 2004). However, unlike most previous implementations, we do not fix the number of shifts or their locations on the phylogeny. Instead, we implement a reversible-jump algorithm that estimates the number, location and magnitude of shifts in adaptive optima, while jointly sampling OU parameters. Although we focus on shifting values of *θ* across the phylogeny, our method can be extended to allow other parameters to differ among regimes (i.e. *α* and *σ*^2^, Beaulieu et al. 2012).

### Reversible-jump Model

We consider a fully resolved phylogeny with *N* taxa. We follow Hansen (1997) and Butler and King (2004) and model a multi-optima OU model with discrete adaptive states ***θ*** = {*θ*_0_, …, *θ*_*K*_}, where *θ*_0_ is the optimum at the root and *K* is the number of shifts between adaptive optima. The locations of shifts between adaptive states is given by a vector of shift locations mapped onto the phylogeny ***L*** = {*L*_1_, …, *L*_*K*_} with each *L*_*i*_ corresponding to beginning of an adaptive regime assigned to the optimum *θ*_*i*_. Given these parameters, the distribution of tip states ***Y*** is multivariate normal with an expectation:

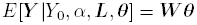

where *Y*_0_ is the root state and ***W*** is a matrix of weights used to calculate the weighted average of adaptive optima, discounted by an exponentially decreasing function that depends on the rate parameter *α* and the elapsed time since the species evolved under a given adaptive optimum (For a full explanation and derivation, see Hansen 1997; Butler and King 2004). The elements of the variance-covariance matrix for ***Y*** are:

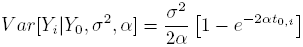

and

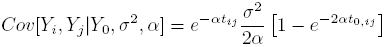

where *t*_0,*i*_ is the time from the root to species *i*, *t*_*ij*_ is the total time separating species *i* and *j*, and *t*_0,*ij*_ is the total time separating the root and the most recent common ancestor of species *i* and *j*.

To calculate the likelihood, we assume that *Y*_0_ = *θ*_0_. Alternatively, the likelihood can be calculated assuming a stationary distribution or by estimating *Y*_0_. However, we found that assuming a stationary distribution resulted in poor mixing when *α* was small (which is typical during the beginning stages of the MCMC). We use a pruning algorithm to speed computation and calculate conditional likelihood (as in FitzJohn 2012). We use a reversible-jump algorithm to search among varying shift numbers (extitK) and shift locations (***L***) (Green 1995; Huelsenbeck et al. 2000, 2004; Eastman et al. 2011; Rabosky et al. 2013). The reversible-jump framework of Green (1995) uses a Metropolis-Hastings algorithm to explore models with varying dimensionality through the course of the MCMC. The amount of time the MCMC spends in a given model is proportional to its posterior probability, thus providing inference on the best supported regime shift placements and magnitudes, while accounting for model uncertainty in all estimated parameters. Proposals in the MCMC are accepted with the general probability:

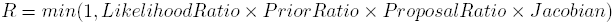

The proposal ratio and the Jacobian together compose the “Hasting’s ratio”, which is dependent on the specific proposals used, as described below.

### Proposals

The proposals to the parameters *α*, *σ*^2^ and *θ*_*i*_ do not change the dimensionality of the model, and can therefore be updated using standard MCMC proposal mechanisms. We update the parameters *α* and *σ*_2_ using a multiplier proposal mechanisms, while *θ*_*i*_ parameters are updated using a sliding-window proposal (described in Eastman et al. 2011).

The dimensionality of the model can change via either a birth or a death step, which adds or subtracts a regime shift from the model, respectively. A branch is chosen from the phylogeny at random. If the branch currently contains a shift, then a death step is proposed. If the branch does not contain a shift, a birth step is proposed. During a birth step, a location for the shift is drawn from a uniform distribution over the length of the branch. The value of the adaptive optima before and after the shift are simultaneously updated according to the following proposal. A random uniform number (*u*) between (−0.5,0.5) is drawn. New values of *θ* are obtained by splitting the value of this random uniform number proportionally to the amount of branch length across the entire tree inherited by each optimum after splitting:

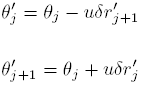

Where 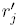 is the proportion of the branch lengths across the tree that will evolve under 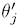 and 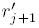 is the proportion of branch lengths that will evolve under 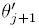, and *δ* is a tuning parameter. Thus, whichever optimum inherits the most branch length will have the most conservative proposal, while the optimum that inherits smaller amounts of branch length will have more liberal proposals (i.e. its value will change the most). The new optimum 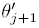 cascades down the phylogeny until it reaches a tip, or a pre-existing shift. The acceptance ratio for the move from parameter set Θ_*K*_ → Θ_*K*+1_ is then (Green 1995; see Online Appendix I for the derivation, which follows Jialin 2012):

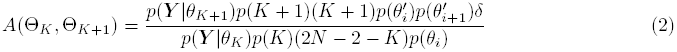

Death steps operate in reverse, collapsing two regimes into a single regime. The new value of this proposed regime is a weighted average of the previous two regimes according to the equation:

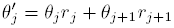

Consequently, proposed values of optima during death steps are deterministic weighted averages, and the acceptance ratio is simply the inverse of equation (2).

In addition to birth and death proposal mechanisms for adding and subtracting regime shifts, we allow shifts to slide along branches without changing the number of parameters in the model with two different proposal mechanisms. The first (*slide*1) allows the position of the shift to move within a branch via a sliding-window proposal mechanism. Proposals are reflected back at the nodes so that the proposed shift location remains on the same branch. A second proposal allows shifts to slide up or down onto neighbouring branches (*slide*2). When this move is proposed, the total number of allowable shifts are counted across the tree for all *K* shifts. Moves are not allowed to branches that already contain shifts, or tipward at the tips and rootward at the root. Each allowable move has an equal probability of being chosen. For example, a shift surrounded by three empty branches (two tipward and one rootward) has three times the probability of being chosen for the proposal than a shift surrounded by only one empty branch and two branches with existing shifts. Once a branch with a shift is chosen, one of the available neighbouring branches is chosen with equal probability. The proposed location of the shift on the new branch is drawn from a uniform distribution and regimes are cascaded tipward until reaching an existing shift.

The proposal probability for the regime birth-death move was set at a fixed value (in our study we used *ϕ*_*bd*_ = 0.45). The remaining proposal probability was divided between the five other proposal mechanisms: the two sliding shift proposals (*slide*1 and *slide*2) and proposals to *α*, *σ*^2^ and ***θ***. We placed equal proposal weight on updating *α* and ***θ***, which were set to 3.5 times the proposal weight of updates to *σ*^2^ and the two sliding shift proposals. We chose these values based on preliminary explorations which indicated that these proposal probabilities resulted in roughly equal effective sample sizes for all parameters. Thus, for 0 < *K* < *K*_*max*_, we set *ϕ*_*α*_ = *ϕ*_***θ***_ = 0.1925 and *ϕ*_*σ*^2^_ = *ϕ*_*slide*1_ = *ϕ*_*slide*2_ = 0.055. When *K* = 0, both *slide*1 and *slide*2 are disallowed, and the proposal probabilities for these two moves are divided evenly between the updates to *α*, *σ*^2^ and ***θ***.

Each shift in our model leads to a unique adaptive optimum. In other words, we do not allow convergence of adaptive regimes in our reversible-jump model (though such models may be fit in 

~~~
bayou
~~~

 with a fixed number of parameters without a reversible-jump proposal). This is because satisfying the “detailed balance condition” of the reversible-jump MCMC requires both forward and reverse proposals to calculate the acceptance probability (Green 1995). However, under convergent regimes, a new shift added to the tree can actually reduce the number of shifts (*K*) by replacing multiple downstream transitions to *θ*_*i*_ in a single move. Since the reverse move would require several proposal steps rather than one, satisfying the detailed balance condition is substantially more complex. It is likely that clever proposal mechanisms can be designed to adequately explore across models with and without convergent regimes with sufficient mixing, but we leave this to future work. Regardless, our primary goal is not to estimate the amount of convergence (which remains quite challenging for single-trait datasets, but see Mahler et al. 2013), but instead to provide a flexible Bayesian framework for exploring among alternative hypotheses and for incorporating prior information to test specific biological interpretations of comparative models. Furthermore, the amount of convergence can be assessed post-hoc by the degree of overlap among regimes, or by comparing the marginal likelihoods among fixed convergent regimes.

### Priors

We placed a prior on the number of shifts between adaptive regimes using a conditional Poisson distribution, with a maximum number of shifts equal to half the number of tips in the phylogeny. Informative priors for this distribution may be taken, for example, from the duration of chronospecies or genera in the fossil record. These may suggest the typical duration of a static adaptive regime. Alternatively, a given adaptive hypothesis may suppose only a few adaptive shifts across a radiation of species.

We placed a normal prior on adaptive optima. Setting a prior on adaptive optima allows us to avoid fitting models in unrealistic regions of parameter space that allow species to track unrealistically distant adaptive optima. For example, if we are studying primate body size evolution, it may be reasonable to take the prior for the adaptive optima of body sizes by using the distribution of body sizes across all terrestrial mammals. Alternatively, we may place the prior based on how distant the optima will be from any extant species. Thus, the prior on the optima would be based on both the observed range of phenotypes in the data being studied plus the prior belief in how distant any given species is from its adaptive optimum. In our model, shifts between adaptive regimes can occur in any region of parameter space. Future implementations could penalize unreasonably large shifts between adaptive regimes by modeling shifts in optima as a compound point process. Landis et al. (2013) proposed fitting models that include these so-called “Levy processes” that model rare jumps in phenotypic space where the rate and magnitude of jumps follow a regular stochastic process model. Integrating Levy models and OU models would add considerable realism to models of phenotypic evolution and allow statistical inference on the rates and distribution of the shifts themselves, but is beyond the scope of the current paper.

Finally, we assign an equal probability of each branch having a shift, with no more than one shift allowed per branch. A more realistic model may allow the probability of shifts between regimes to occur proportional to the length of the branch, and to allow multiple shifts per branch. We leave these enhancements to future work. Shifts may occur anywhere on a branch with a uniform probability distribution, thereby allowing uncertainty in the location of the shift to affect estimation of other parameters.

### Simulation study

To assess the performance of the method given the assumptions we have made above, we conducted a simulation study. Phylogenies were simulated under a pure-birth process using the R package TreeSim (Stadler 2011) with 64 tips (except for when the effect of sample size was being examined, see below) and a constant birth rate of 0.1. For all simulations, the resulting phylogenies were scaled to unit height and starting parameters were drawn from the prior distribution to initialize the chain. Two independent chains were run for at least 200,000 generations, with a thinning interval of 20 and the first 60,000 generations discarded as burnin. We estimated Gelman’s R-statistic for each parameter, which compares the within- to between-chain variance to evaluate convergence (Gelman and Rubin 1992). Values of R close to 1 indicate that the two chains are not distinguishable, whereas high values (we used a cutoff of R = 1.1) indicate non-stationarity of the chains.

We tested the performance of the method across a range of parameter values varying a single parameter at a time and using broad, weakly informative priors. For the standard simulation we used *α*=3 (phylogenetic half-life = 23.1% of the total tree height), *σ*^2^ = 3 (stationary variance = 0.5), *K* = 9 (10 selective optima including the root) and a root value of 0. Shifts were randomly placed on available branches at regular intervals separated by 0.1 units of tree height. At each shift, new optima were drawn from a normal distribution with mean = 0 and sd = 3. Each parameter was varied independently for a series of simulations while all others were held constant at the above values. Parameters were examined over the following ranges: Phylogenetic half-life (*ln*(2)*/α*; in units of tree height) = (0.01, 0.05, 0.1, 0.2, 0.3, 0.4, 0.5, 0.75, 1, 2, 10), *σ*^2^ = (0.1, 0.5, 1, 2, 3, 5, 10, 25, 50), clade size = (32, 64, 128, 256, 512, 1064) and *K* = (1, 5, 7, 8, 9, 10, 11, 13, 20, 50).

We used the simulation-based posterior quantile test of Cook et al. (2006) to validate our implementation. This powerful method relies on the distributional properties of Bayesian posteriors when simulation parameters are drawn from the prior distribution. Specifically, if the software and model are correctly formulated, then posterior quantiles should contain the true parameter value in the corresponding percentage of simulations (e.g. a 50% posterior quantile will contain the true parameter value in 50% of the simulations). Although validation using the posterior quantile test does not guarantee that an implementation is correct, it is a highly sensitive method of testing the null hypothesis that the software is working correctly (Cook et al. 2006). Parameters were simulated from the prior distribution, which were set as follows: P(*α*)∼ LogNormal(ln *μ*=0.25,ln *σ*=1.5); P(*σ*^2^) ∼ LogNormal(ln *μ*=0.25,ln *σ*=0.1); P(*θ*_*i*_) ∼ Normal(*μ* = 0, *σ* = 3); P(*K*) ∼ Conditional Poisson(λ = 10, *K*_*max*_ = 32). Data were simulated for each set of parameters on a simulated phylogeny (following the same simulation parameters described above) and posterior distributions were estimated. The quantile of the true parameters are determined within the estimated posterior distribution. If the posterior distribution is estimated correctly, then the distribution of these quantiles across simulations should follow a uniform distribution (Cook et al. 2006). We tested each of the parameters *α*, *σ*^2^, *K* and *θ*_0_ (the root optimum) for deviations from a uniform distribution using a Kolmogorov-Smirinov test. Significant deviations would indicate that the method incorrectly estimates the posterior distribution.

We used these same simulations from the posterior quantile test (in which simulation parameters were drawn from the prior distribution) to evaluate the performance of the method in assessing the location of shifts and magnitude of shifts. Thus, we assessed the power of the method to detect shifts across a broad range of possible parameter combinations. Posterior probabilities were calculated for each branch by counting the proportion of posterior samples with a shift on that branch, after excluding the burn-in phase. Consequently, a posterior probability of 0.5 would indicate that half the posterior sample contained that shift on a given branch. We then plotted the posterior probability of each branch against two values. The first we term “regime divergence” and is defined as the log of the ratio between the shift magnitude and the stationary variance of the OU process. The second we term the “scaled age” of the shift, which is the log of the ratio between the age of the shift (in time units before present) and the true phylogenetic half-life value. We expect a relationship between these two ratios and our power to detect shifts because 1) higher values of regime divergence indicate a more dramatic shift relative to the variation within a selective regime and 2) the scaled age is a measure of how recent the shift is relative to the speed of the OU process. A very low scaled age value would indicate that very little time has elapsed for the descendants of this shift to adapt to the new regime, thus resulting in low power to detect the shift. Conversely, high values would indicate that ample time has elapsed for species to equilibrate on their new adaptive optimum. We estimated a contour plot of the posterior probability values for all branches across simulation against regime divergence and scaled age by kriging. In addition to the above analyses, we conducted a number of additional simulations to assess the power of the method to accurately recover shift number, magnitude and location, including an extensive prior sensitivity analysis for the prior on the number of shifts. A detailed description of these simulations can be found in the Online Appendix III.

### Testing specific biological interpretations of evolution

One of the strengths of OU models is that the process and parameters describe the dynamics of adaptive evolution in biologically interpretable quantities. However, whether or not we can interpret OU models fit to the deep timescales of phylogenetic comparative data as–for example–stabilizing selection and genetic drift, has generally been addressed with only qualitative arguments. The utility (and in some cases drawback) of the Bayesian approach is that it allows/requires the use of priors on parameter values. We use this feature in 

~~~
bayou
~~~

 to allow the specification of informative priors on parameter values that correspond to particular biological interpretations of the OU process. For example, rather than specifying a prior on *α* and *σ*^2^, we may instead have prior information on phylogenetic half-life or stationary variance. Reparameterization of the model in this way would provide a useful way to incorporate prior information on the width of niches, adaptive peaks, adaptive zones or the rate of adaptation toward a phenotypic optimum.

We use the quantitative genetic model of Lande (1976), who showed that genetic drift around a stationary Gaussian adaptive peak results in a OU process with parameters:

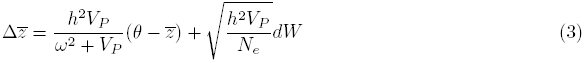

where *h*^2^ is the trait heritability, *V*_*P*_ is the phenotypic variance, *ω*^2^ is the width of the adaptive landscape, and *N*_*e*_ is the variance effective population size. We allow specification of this model in 

~~~
bayou
~~~

 and use informative priors on these parameters to constrain the model to realistically reflect this particular biological interpretation. Note that the branch lengths of the phylogeny should be expressed in number of generations in order to fit such a model. Furthermore, we assume that all parameters are constant across the phylogeny, an assumption that is likely to be violated.

To test the utility of this method, we compared models fit using either an unconstrained standard OU parameterization (*OU*_*Free*_) or a quantitative genetic Lande model parameterization (*QG*) with priors taken from compilations of empirical estimates typical for linear body size traits on the log scale [Table 1; similar to the approach of Estes and Arnold (2007), but in an explicitly Bayesian framework]. While the priors for the *OU*_*Free*_ model were chosen to have a majority of the prior density centered on values of parameters typical of comparative data, effort was made to make them broad enough to also have a some significant prior density on the values generated under Lande-model priors. As before, we simulated 64-taxa trees, but scaled trees to 50 million years old and a generation time of 5 years. Simulated data were drawn by drawing parameter values from either the *QG* or *OU*_*Free*_ prior distributions. We included normally distributed measurement error in both the simulation and estimation of the data, with a error variance of 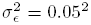.

**Table 1.** Prior distributions for *OU*_*Free*_ and *QG* models used in simulation study modeling different biological processes. Models are simulated and fit to the data from these priors, intended to reflect values typical for linear ln(body size) related measurements in a clade 50 my old that spans two magnitudes in size. Measurement error (ME) is simulated and fit to be ∼Norm (μ = 0, 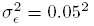.

| Parameter | Prior | Quantiles (1%, 50%, 99%) | Ref |
| --- | --- | --- | --- |
| **Model: $QG$** |  |  |  |
| $h^2$ | $\sim \text{Beta}(a = 15, b = 20)$ | (0.25; 0.43; 0.62) | Mousseau et al. 1987 |
| $V_P$ | $\sim \text{LogNorm}(ln\mu = -5, ln\sigma = 0.8)$ | (0.032<sup>2</sup>; 0.082<sup>2</sup>; 0.208<sup>2</sup>) | Uyeda et al. 2011 |
| $\omega^2/V_p$ | $\sim \text{LogNorm}(ln\mu = 4, ln\sigma = 1.5)$ | (1.67; 54.6; 1,790) | Estes and Arnold 2007 |
| $N_e$ | $\sim \text{LogNorm}(ln\mu = 10, ln\sigma = 2)$ | (210; 22,000; 2.31E6) | Estes and Arnold 2007 |
| Phy half-life (y) | - | (21.2; 459; 15,200) |  |
| $\sqrt{V_y + \sigma_\epsilon^2}$ | - | (0.05; 0.05; 0.079) | |
| **Model: $OU_{Free}$** |  |  |  |
| $\alpha$ | $\sim \text{LogNorm}(ln\mu = -3, ln\sigma = 3)$ | (4.6E-5; 5.0E-2; 53) | Hansen 2012* |
| $\sigma^2$ | $\sim \text{LogNorm}(ln\mu = 0, ln\sigma = 2)$ | (9.5E-3; 1; 105) | |
| Phy half-life (my) | - | (0.013; 13.9; 1,487) |  |
| $\sqrt{V_y + \sigma_\epsilon^2}$ | - | (0.069; 3.17; 210) | |
| **Both models** |  |  |  |
| $\theta$ | $\sim \text{Norm}(ln\mu = 0, ln\sigma = 0.5)$ | (-1.2; 0; 1.2) | |
| $K$ | $\sim \text{CondPoisson}(\lambda = 15, K_{max} = 32)$ | (7; 15; 25) | |

\*This prior on $\alpha$ puts about $\sim 20\%$ of the probability density on a phylogenetic half-life less than the average youngest splits in the simulated phylogenies ( $\sim 1$ my, white noise-like evolution) and $\sim 30\%$ of the probability density above the tree height (50 my, i.e. BM-like evolution).

To compare model fits to the simulated data, we estimated marginal likelihoods under each model using stepping-stone sampling of models fit to either the *OU*_*Free*_ or *QG* parameterizations (Fan et al. 2011). These marginal likelihoods were then used to compute Bayes Factors for model selection (Kass and Raftery 1995). Briefly, the stepping-stone method runs a sequence of MCMC simulations to estimate “power posterior” distributions sequentially stepping from a reference distribution (we used uncorrelated multivariate distributions with parameters estimated from the posterior distribution) to the posterior distribution (Fan et al. 2011). Each power posterior is used as an importance density to estimate a series of ratios of normalizing constants, the product of which provides the ratio of the known reference distribution and the marginal likelihood (Xie et al. 2011). We used a total of 12 steps along the path from the reference distribution to the posterior for each stepping-stone MCMC and ran each step for 500,000 generations. This was done for 10 datasets simulated from the prior for each model. Distributions of Bayes Factors were then compared across model fits to evaluate the degree of support for a specific biological interpretation of the model under different simulation parameters.

### Test Case: Chelonian carapace evolution

The wide variation in body sizes across extant chelonians (tortoises and turtles) has been hypothesized to result from a number of causes, including gigantism resulting from marine and island habitats that are thought to remove evolutionary constraints on body size (Arnold 1979). Jaffe et al. (2011) explored the evolution of the length of the carapace (i.e. the dorsal shell) in a phylogeny of 226 extant chelonians using traditional likelihood approaches by fitting multi-optima OU models to the data using 

~~~
ouch
~~~

 (Butler and King 2004). The best-fitting model in Jaffe et al. (2011) assigned four separate regimes to freshwater, mainland-terrestrial, island-terrestrial, and marine species of turtles (which they called the ’OU2’ model; we will refer to the model as the *OU*_*habitat*_ model). This model was compared to a limited set of alternative hypotheses, including BM, a single-optimum OU model, and several models which collapsed various combinations of the four regimes in the *OU*_*habitat*_ model. From these results, Jaffe et al. (2011) concluded that there was a strong signal in the data supporting a shift to larger size optima in chelonians in marine and island habitats. Parameter estimates from the study suggest that body size evolves slowly toward these new optima, with phylogenetic half-lives on the order of 15-20 million years (7-10% of tree height). This same dataset served as a test case for a reversible-jump algorithm that explored the potential for shifts in BM rate parameters (Eastman et al. 2011). The relaxed BM model (rBM) uses a reversible-jump framework to find shifts in evolutionary rates in a manner very similar to 

~~~
bayou
~~~

. For the Jaffe dataset, Eastman et al. (2011) found shifts in evolutionary rates between several groups of turtles and tortoises, including increased rates in Testudinidae (tortoises) and Emydidae (pond turtles). Note that we discovered an error in the acceptance ratio of the rBM model in previous versions of the R package 

~~~
auteur
~~~

 (Eastman et al. 2011). We provide the corrected acceptance ratio for the rBM model in Online Appendix II, which has been updated for the implementation of the rBM model in the R package 

~~~
geiger
~~~

 (versions 1.99 and greater, Harmon et al. 2008). Because the Eastman et al. (2011) model detects rate-shifts not mean-shifts, it is not well-suited for testing hypotheses about directional adaptation in a clade to an optimal state. We instead fit a related, recently developed model of phenotypic shifts in a BM framework that combines “jumps” or mean-shifts with a standard BM (bm-jump model, Eastman et al. 2013). Note that this model is distinct from the models implemented in 

~~~
bayou
~~~

, because mean-shifts under the bm-jump model come from a distinct distribution and are best thought of as temporary shifts to high rates of evolution. By contrast, OU models use the same *α* parameter to control the rate of adaptation to a new optimum before and after a shift, and therefore cannot combine rapid jumps in mean with BM-like evolution.

We use the Jaffe et al. (2011) dataset to demonstrate the utility of 

~~~
bayou
~~~

, and to compare to existing approaches. We analyzed the *OU*_*habitat*_ model in 

~~~
bayou
~~~

 by constraining the location of shifts and regimes to be the same as the model of Jaffe et al. (2011), except we allowed the location of the shift to move freely along the branch (rather than constraining it to occur at the nodes). We replicated maximum likelihood estimates for the *OU*_*habitat*_ model to obtain comparable estimates of parameters using the R package 

~~~
OUwie
~~~

 (Beaulieu et al. 2012). We then compared this to an unconstrained 

~~~
bayou
~~~

 model that allows shifts to be assigned freely among the various branches. Note that these models are not nested, because in the *OU*_*habitat*_ model regimes are convergent, whereas in the unconstrained model, each shift is given its own unique adaptive regime. Priors on the parameters in all runs were assigned as follows: P(*α*)∼ LogNormal(ln *μ*=-5,ln *σ*=2.5); P(*σ*^2^) ∼ LogNormal(ln *μ*=0,ln *σ*=2); P(*θ*_*i*_) ∼ Normal(*μ* = 3.5, *σ* = 1.5); P(*k*) ∼ Conditional Poisson(*λ* = 15, *k*_*max*_ = 113).

We compared these models by comparing the posterior probabilities of shifts on each branch in the 

~~~
bayou
~~~

 unconstrained run to the locations of the shifts in the *OU*_*habitat*_ hypothesis. Furthermore, we evaluated overall model support for the constrained vs. the unconstrained model by using stepping-stone sampling using the method of Fan et al. (2011) to estimate Bayes Factors. Note that Bayes Factors can be sensitive to prior specification, especially when one model has a very specific prior specification (e.g. the location and number of shifts are fixed, as in the *OU*_*habitat*_ model) and the alternative has a very vague prior (as in the unconstrained 

~~~
bayou
~~~

 model). Specifically, vague priors are expected to produce lower marginal likelihoods than more specific priors, and thus these tests favor constrained models. Furthermore, the *OU*_*habitat*_ model has fewer parameters than the unconstrained 

~~~
bayou
~~~

 model with the same number of shifts due to evolutionary convergence.

In addition to comparing the unconstrained and constrained model in a Bayesian framework, we compared our method to the method of Ingram and Mahler (2013) as implemented in the R package 

~~~
SURFACE
~~~

. Our unconstrained model is analogous to the forward-addition procedure of 

~~~
SURFACE
~~~

, and thus we compare parameter estimates and shift locations in both the forward and reverse steps of the 

~~~
SURFACE
~~~

 algorithm independently. These two methods differ in that the 

~~~
SURFACE
~~~

 algorithm of Ingram and Mahler (2013) relies on stepwise-AIC rather than Bayesian reversible-jump methods to find the location of shifts. While stepwise-AIC methods attempt to find the single, best-fitting model, our method integrates over uncertainty in regime placement and returns a posterior of shift locations that well-describe the data. Finally, we fit the BM-jump model of *auteur*, which fits jumps to a background of constant-rate BM (Eastman et al. 2013). As we described above, this is similar to the 

~~~
bayou
~~~

 model, but differs in that jumps are realized instantaneously rather than approaching at a rate determined by *α*. Thus, these two models may be expected to find similar shift locations and have relatively comparable parameter estimates. As with the 

~~~
bayou
~~~

 implementation, we place a conditional Poisson prior on the number of shifts (“jumps”) at λ = 15.

## Results

### Simulation study

Under most simulation conditions, convergence was reached within 500,000 generations (i.e. Gelman and Rubin’s R reached below 1.1 for all parameter values). When convergence failed to occur within this time frame, it tended to be when shift magnitudes were large relative to the stationary variance of the process (i.e. high regime divergence). This results in steep likelihood peaks and a tendency to get trapped under non-optimal configurations of shifts. A potential solution to this issue, besides running longer and more chains, would be to use Metropolis-Coupled MCMC, which improves mixing by implementing “heated” chains that explore the likelihood surface more efficiently. Interestingly, these instances in which mixing is poorest and convergence is most problematic are the instances in which the model finds the strongest support for regime shifts and estimates their location most reliably (although it may get caught in less parsimonious likelihood peaks than the simulated model). Nonetheless, the model effectively identifies the presence of adaptive peaks shifts, although it may have difficulty determining the exact order of shifts among branches in these instances.

Overall, simulations indicated that parameters are estimated with reasonable accuracy. The diffusion rate parameter *σ*^2^ tends to be slightly underestimated, especially for low phylogenetic half-life values (i.e. values less than most of the splits in the tree) and overestimated when phylogenetic half-lives are much longer than tree height (Figure 1, A & D). However, for half-life values ranging from around 0.1-2 times tree height, *σ*^2^ is estimated reasonably with a slight bias toward underestimation (Figure 1, D). Phylogenetic half-life tends to be overestimated over most of this range as well (Figure 1, B & E). These two effects balance out, however, and produce reasonable estimates of stationary variance. Increasing number of tips greatly improves estimates of both *σ*^2^ and *α* and reduces bias (Figure 1, J & K). For small numbers of tips (e.g. 32), there is a tendency to fit models with higher values of phylogenetic half-life (more BM-like) models. This is likely because many shifts occur on branches leading to singletons, resulting in very little power to distinguish the effect of rate parameters (*α* and *σ*^2^) from shifts (*θ*).

We show that inference becomes problematic when the ratio of the number of tips to the number of shifts decreases below 4, and recommend at a minimum ∼ 50 tips. The number of estimated shifts seems are particularly sensitive to the prior when using the conditional Poisson as the prior, despite some influence of the data on the estimation of the number of shifts (Figure 1, C, F, I & L; Online Appendix III). When the number of shifts are large, the method generally results in underestimation of the true number of shifts. This is particularly true when more permissive priors are used (e.g. negative binomial or discrete uniform, Online Appendix III). However, even when the number of shifts is not reliably estimated, parameter estimates for *α*, *σ*^2^ and branch-specific posterior probabilities were not substantially affected until the number of shifts was large (e.g. 20 shifts on a 64-128 taxa phylogeny, Figure 1, G-I; Online Appendix III).

**Figure 1.**
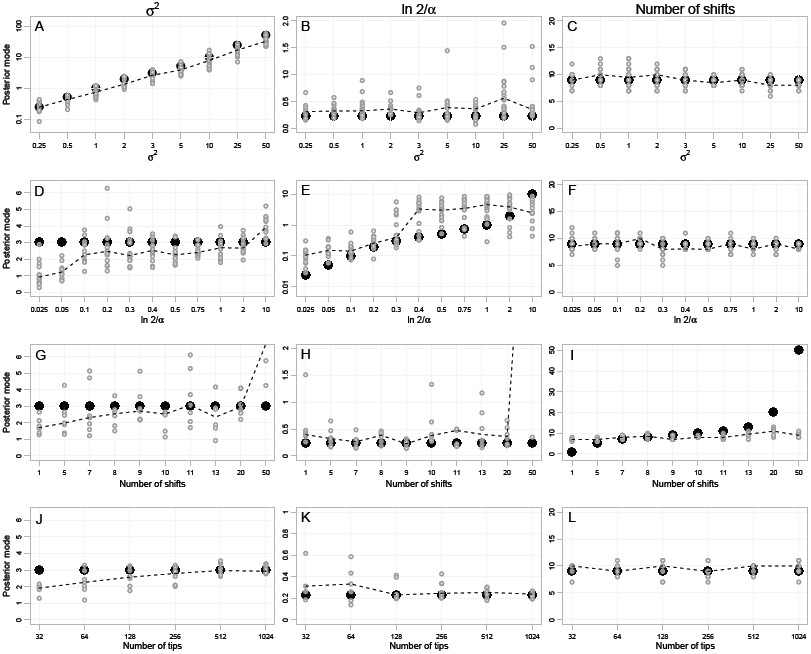
Results of simulation study varying the parameters *σ*^2^ (row 1, A-C), phylogenetic half-life (*ln*(2)/*α*; row 2, D-F), the number of shifts on the phylogeny (*K*; row 3, G-I) and the number of tips in the phylogeny (row 4, J-L). Solid black points indicate the true value used to simulate the data, dotted lines indicate the median posterior mode across simulations. Priors for parameters are as follows: *α* ∼ LogNormal(ln *μ* = 0.25, ln *σ* = 1.5), *σ*^2^ ∼ LogNormal(ln *μ* = 0, ln *σ* = 5), *θ* ∼ Normal(*μ* = 0, *σ* = 3), *K* ∼ Conditional Poisson (λ = 9, *K*_*max*_ = ntips / 2).

Inferences based on branch-specific posterior probabilities were not affected by mis-estimation of the total shift number, as long as the prior allowed a sufficient number of shifts to explain the true model (Online Appendix III). Rather than producing a large number of strongly supported false-positives, strong priors with a large number of expected shifts (e.g. conditional Poisson with λ = 20) resulted in diffusely elevated branch posterior probabilities across all branches when no high magnitude shifts were present. By contrast, when the prior allows only a few shifts, complex models are poorly fit and the models tend to collapse to BM-like models with mis-estimated parameters. Thus, priors that favor complex models have little effect on inference of shift location and magnitude, whereas conservative priors tend only to fit models with very few shifts, and severely mis-estimate parameters for complex models (see Online Appendix III).

Branches on which a shift was simulated have much higher posterior probabilities than branches in which no shift occurred in the true simulation model, even when the number of shifts is over or under-estimated (Figure 2). This is particularly true for small *σ*^2^ and low phylogenetic half-life (Figure 2A & B). This is because low values of *σ*^2^ results in very little trait variation within regimes relative to the divergence between regimes (high regime divergence), and low values of phylogenetic half-life result in rapid adaptation to regimes (high scaled age). As phylogenetic half-life approaches 1 (i.e. the tree height), the power to detect shifts is reduced nearly to 0 under the simulation parameters we examined. The number of tips did not significantly affect the posterior probability of correctly identifying a shift (Figure 2D). This is likely because the expected number of tips evolving under a randomly placed adaptive regime increases slowly with increasing number of tips, meaning that adding tips does relatively little to improve estimates of shift parameters (i.e. assuming equiprobable shift locations among branches–regardless of the phylogeny size–results in approximately 50% of randomly placed shifts being placed on terminal branches that contain only one descendant, Ho and Ané 2013). Furthermore, as the number of tips increased, the prior probability of a given branch having a shift decreased. When errors were made and branches with a true shift were assigned low posterior probabilities, this was generally because the shift was assigned with high posterior probability to a neighbouring branch.

**Figure 2.**
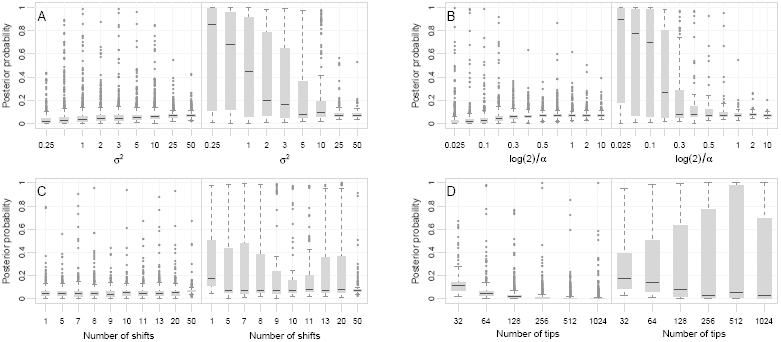
Estimation of branch posterior probabilities using the same simulations as in Figure 1 for varying values of (A) *σ*^2^, (B) phylogenetic half-life, (C) number of shifts (*K*) and (D) number of tips. For each plot, boxplots indicate the distribution of posterior probabilities of a shift occurring on branches that either contain no shift in the true model (left side of each panel) or contain a shift (right side). Locations of shifts were chosen randomly across the phylogeny, and magnitudes were determined by optima drawn randomly from a normal distribution.

Simulation of parameters from the prior distribution and subsequent simulation was carried out for 1118 simulations run for 200,000 generations to evaluate posterior quantiles of the simulated parameter values (Cook et al. 2006). A total of 938 of the simulation runs resulted in Gelman’s R statistics less than 1.1 and were used in subsequent analyses. Posterior quantiles of the simulated parameter values for the root (*θ*_0_), the number of shifts (*K*) and *σ*^2^ were not significantly different from a uniform distribution, as expected if these parameters are estimated accurately and without bias. Posterior quantiles for the *α* parameter deviated significantly from a uniform distribution (p = 0.01, Table 2), tending to be slightly under-estimated (∼ 54% of true parameter values were estimated to be in the upper 50% quantile of the posterior distribution). This significant deviation may have been affected by issues with convergence rather than implementation error, as *α* tended to converge slower than other parameters and runs that failed to converge (∼ 16.1% of simulation runs) had significantly larger values of *α* (*p* < 0.05), suggesting that systematic removal of non-convergent runs could affect the distribution of posterior quantiles. Regardless, the distribution for *α* was nonetheless qualitatively quite similar to a uniform (Figure 4). Overall, the method appears to perform quite well at recovering accurate posterior distributions for the estimated parameters.

**Table 2.** Posterior quantiles for parameter values and p-values from a Komolgorov-Smirnov test. Runs that did not converge were removed from the analysis.

|  | N | 2.5% | 25% | 50% | 75% | 97.5% | p-value |
| --- | --- | --- | --- | --- | --- | --- | --- |
| Log Likelihood | 938 | 0.03 | 0.26 | 0.53 | 0.76 | 0.98 | 0.17 |
| Log Prior | 938 | 0.02 | 0.30 | 0.57 | 0.80 | 0.98 | 0.00 |
| $\alpha$ | 938 | 0.02 | 0.26 | 0.54 | 0.78 | 0.99 | 0.01 |
| $\sigma^2$ | 938 | 0.02 | 0.26 | 0.52 | 0.74 | 0.98 | 0.78 |
| $K$ | 938 | 0.03 | 0.27 | 0.51 | 0.76 | 0.99 | 0.42 |
| Root $\theta_0$ | 938 | 0.02 | 0.22 | 0.51 | 0.76 | 0.98 | 0.29 |

**Figure 4.**
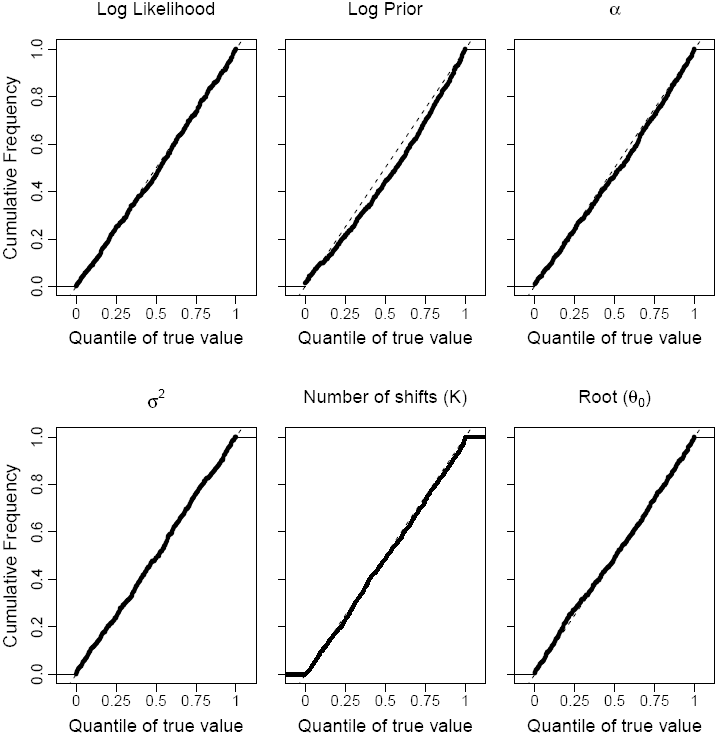
Cumulative distribution plots for posterior quantiles for selected parameters. If posteriors are estimated accurately, then the quantiles of true values of the parameters across simulated datasets should be uniformly distributed (Cook et al. 2006) and follow the dotted lines, which indicate the expected cumulative distribution function for a uniform distribution. Each MCMC was run for 200,000 generations and the first 30% of the samples were discarded as burnin. Runs in which Gelman’s R failed to reach below 1.1 at the end of the run were removed. A total of 938 simulations were used after removing runs that did not reach convergence.

Branches with shifts tend to have high posterior probabilities if the shift magnitude is high relative to the stationary variance and when the age of the shift is much older than the phylogenetic half-life (Figure 3). Specifically, a shift that is 1 phylogenetic half-life old and 1 standard deviation from the stationary distribution away from the previous optimum will have an estimated posterior probability of having a shift of around 0.2, or slightly more than twice the prior probability of a branch having a shift (Figure 3).

**Figure 3.**
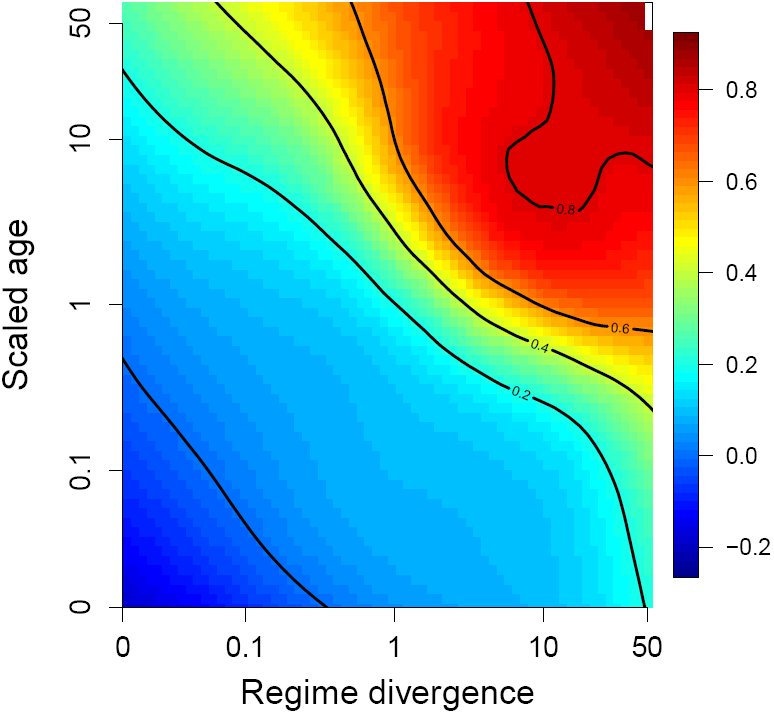
Relationship between regime divergence 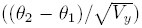, the scaled age of the shift (*shift age/phylogenetic half-life)* and the posterior probability of detecting a shift. All branches from each 64-taxon tree were plotted to estimate this Surface using ordinary kriging to visualize the relationship. Prior probability of a branch being selected is 0.0827. Branches without shifts were given a small divergence value (log(0.01)) and correspond to the left-most data in the plot.

### Modeling biological interpretations

Models simulated under Lande model (*QG*) priors were decisively favored under a *QG* parameterization as opposed to an *OU*_*Free*_ in 9 out of 10 simulations, with exceedingly high Bayes Factors (Figure 5, mean = 4746.5, sd = 6166.4). Around half of these simulations failed to converge across chains (Gelman and Rubin’s R > 1.1) after 500,000 generations, and therefore a total of 20 simulations were required to obtain 10 well-estimated Bayes factors. Models simulated under *OU*_*Free*_ priors typical of comparative data also resulted in decisive support in all 10 simulations against the Lande model, but Bayes Factors were not nearly as high as those simulated under the Lande model (Figure 5, mean = 46.6, sd = 15.7). The asymmetry results from the known effect of vague priors on Bayes Factors (Kass 1993) when compared to highly specific models. Even so, we observe that we can easily reject certain parameterizations and prior sets over alternatives.

**Figure 5.**
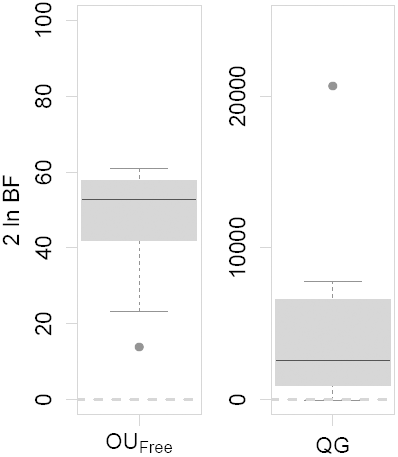
Model comparison for biological interpretations of OU models. Models were simulated either under diffuse priors typical of comparative data (*OU*_*Free*_, left) or under realistic priors for the Lande model (*QG*, right). Both *OU*_*Free*_ and *QG* models were then fit to the data, and marginal likelihoods were estimated using stepping-stone modeling to obtain Bayes Factors (BF). 2 ln BF are shown, with values above 0 (dotted gray line) indicating that the true model was favored. A total of 10 simulations were run under each model (see text for details).

### Chelonia carapace evolution

The unconstrained 

~~~
bayou
~~~

 model identified a number of highly supported shifts in the posterior distribution (Figure 6). In particular, strong shifts were detected for increased carapace length in the clades that include softshell turtles (family Trionychidae), sea turtles (superfamily Chelonioidea), the genus *Batagur*, the Malaysian giant turtle (*Orlitia borneensis*) and two clades of tortoises (Figure 6). Decreases in size were found in the clade leading to tortoises and most modern turtles, as well as a few moderately support shifts across the tree, such as the tortoise clade including *Geochelone elegans* and *Geochelone platynota*. Posterior distributions of parameter values were much narrower than the prior distributions, and indicate substantial information in the data driving the estimation of these parameters. Phylogenetic half-lives were relatively short compared to the height of the tree (posterior median = 3.94 my) indicating that after accounting for adaptive regimes, very little phylogenetic covariance remained among species in the phylogeny. This is in contrast to the *OU*_*habitat*_ model, which estimated significantly higher values of phylogenetic half-life and lower values of *σ*^2^ (Figure 7).

**Figure 6.**
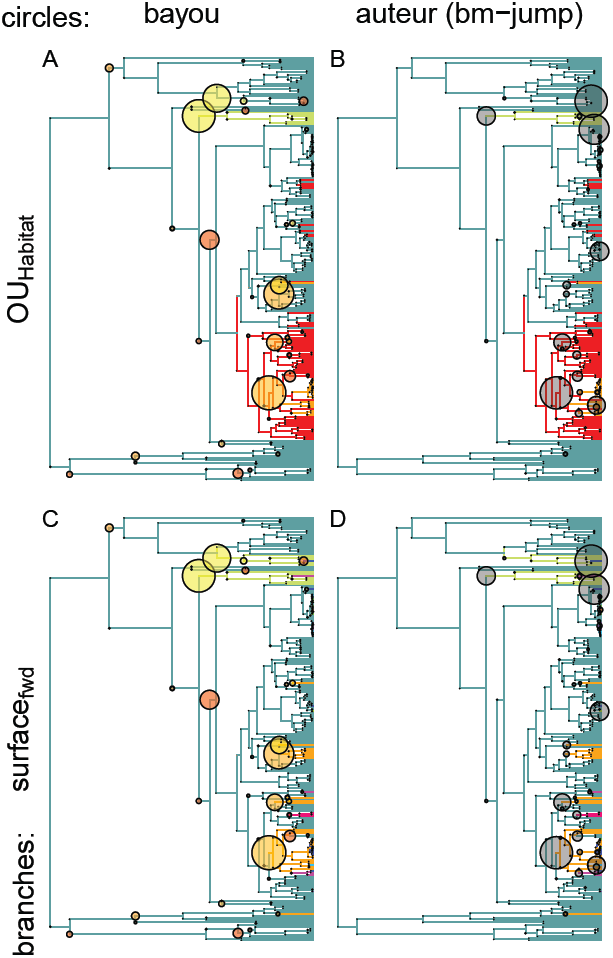
Model fits of multi-optima OU models using different methods. Circles at the nodes in A) and C) indicate by their size the posterior probability of shift locations from the reversible-jump model implemented in 

~~~
bayou
~~~

, with the larger phenotypic values for optima being indicated by yellow and smaller optima indicated by red. Circles in B) and D) are the posterior probability of a shift in a constant-rate BM model with jumps fit using the bm-jump model in 

~~~
auteur
~~~

. Branch painting in A) and B) is the *OU*_*habitat*_ model (which corresponds to the OU2 model of Jaffe et al. 2011). Regime paintings in C) and D) correspond to the best fitting model from 

~~~
SURFACE
~~~

 using forward stepwise addition (Ingram and Mahler 2013).

**Figure 7.**
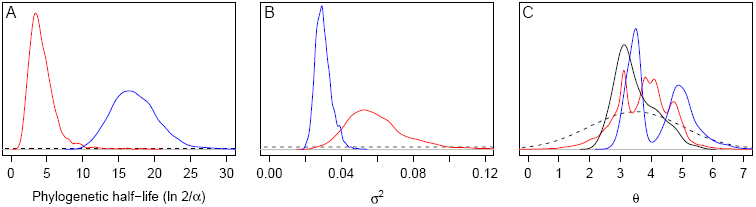
Posterior distributions for parameters estimated using 

~~~
bayou
~~~

 under a reversible-jump (Red) and fixed *OU*_*habitat*_ (Blue) models to the chelonia carapace data. Parameters estimated include A) phylogenetic half-life B) *σ*^2^ and C) the distribution of phenotypic optima (*θ*). Dotted lines indicate prior density, black solid line in C) indicates the distribution of phenotypes in the data.

Support for the unconstrained 

~~~
bayou
~~~

 model over the Bayesian *OU*_*habitat*_ model was very strong, with a 2 ln BF = 15.24 (Kass and Raftery 1995,; Table 3). Only one shift (leading to sea turtles) identified in the *OU*_*habitat*_ model was identified as strongly supported in the posterior of the unconstrained 

~~~
bayou
~~~

 model (Figure 6), while 

~~~
SURFACE
~~~

 and 

~~~
auteur’s
~~~

 bm-jump model found more comparable shift locations. We conclude that while the *OU*_*habitat*_ model is better than many models, it is not a representative model from the posterior distribution obtained from 

~~~
bayou
~~~

. Instead, the hypothesis proposed by Jaffe et al. (2011) captures only a few of the relevant statistical features of the data. The *OU*_*habitat*_ model also had higher estimates for both the phylogenetic half-life and for stationary variance, likely because multiple adaptive regimes were combined into single adaptive regimes, resulting in inference of broader adaptive zones, and a weaker rate of adaptation.

**Table 3.**
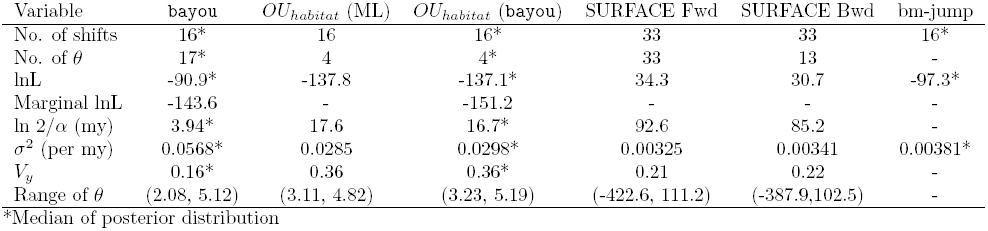
Comparison of model fits to Chelonia carapace length data (Jaffe et al. 2011). The range of *θ* values are taken from either the posterior distribution (for Bayesian analyses) or from the range of estimates for individual optima from the ML analyses.

| Variable | bayou | $OU_{habitat}$ (ML) | $OU_{habitat}$ (bayou) | SURFACE Fwd | SURFACE Bwd | bm-jump |
| --- | --- | --- | --- | --- | --- | --- |
| No. of shifts | 16* | 16 | 16* | 33 | 33 | 16* |
| No. of $\theta$ | 17* | 4 | 4* | 33 | 13 | - |
| lnL | -90.9* | -137.8 | -137.1* | 34.3 | 30.7 | -97.3* |
| Marginal lnL | -143.6 | - | -151.2 | - | - | - |
| $\ln 2/\alpha$ (my) | 3.94* | 17.6 | 16.7* | 92.6 | 85.2 | - |
| $\sigma^2$ (per my) | 0.0568* | 0.0285 | 0.0298* | 0.00325 | 0.00341 | 0.00381* |
| $V_y$ | 0.16* | 0.36 | 0.36* | 0.21 | 0.22 | - |
| Range of $\theta$ | (2.08, 5.12) | (3.11, 4.82) | (3.23, 5.19) | (-422.6, 111.2) | (-387.9, 102.5) | - |

\*Median of posterior distribution

~~~
SURFACE
~~~

 runs identified most of the same shifts that were found in 

~~~
bayou
~~~

, as well as considerable amounts of convergence. However, more shifts were identified, likely due to the prior on the number of shifts (

~~~
SURFACE
~~~

 identified 33 shifts, while the prior on the number of shifts in the 

~~~
bayou
~~~

 runs was only 15). Estimates of adaptive optima, ***θ***, were extremely distant in the best-fitting 

~~~
SURFACE
~~~

 models, and estimates of phylogenetic half-life were correspondingly considerably larger than the estimates from bayou (Table 3).

## Discussion

### Bayesian Inference of Adaptive Regimes

In this study, we have shown how Bayesian inference of adaptive regimes fit to multi-optima OU models provides a flexible framework for testing evolutionary hypotheses. Bayesian OU models have had limited application to trait evolutionary studies (but see Reitan et al. 2012, for layered OU models applied to fossil timeseries data), but offer a number of distinct advantages over existing methods.

First, our method integrates over uncertainty in regime placement and allows inference of the location, magnitude and number of adaptive shifts. Based on our simulation results we find that inference on the number of shifts is more difficult as estimation of this parameter tends to heavily influenced by the prior distribution. However, other parameters in the model are well-estimated and the method correctly identifies the location of most shifts in the phylogeny so long as the number of shifts is not large (*K* > 25% the number of tips). The probability of correctly identifying a shift increases with the magnitude and age of adaptive regimes. Because low magnitude, recent shifts of little effect can always be added to the model, inference should focus on the branch posterior probabilities themselves rather than the total shift number (see Online Appendix III).

Second, the great advantage of OU models is their compatibility with our understanding of the evolutionary process (Hansen 1997; Hansen et al. 2008). On the other hand, many of the statistical properties of OU models can complicate inference resulting from inconsistent estimators and lack of identifiability (which has been shown for the case when the root state is drawn from a stationary distribution, see Ho and Ané 2013). We show how Bayesian implementation of these models allows a full exploitation of the biological realism of OU models by allowing the use of informative priors that constrain the model to biologically realistic values while simultaneously alleviating the some of the statistical issues of OU model-fitting, such as the existence of likelihood ridges that extend into regions of unrealistic parameter space. Furthermore, we demonstrate how the model can be used to explicitly test alternative biological interpretations by testing alternative parameterizations of the models. Note that no method, however, will retrieve an automatic determination of whether a model is correct or not, only whether the correlative pattern and model is consistent with a given biological interpretation. The discretion of the researcher is needed to adequately interpret the reasonableness of the results produced by 

~~~
bayou
~~~

.

Just as a clear understanding of the mechanisms behind molecular evolution have revolutionized methods of phylogenetic inference (Kimura 1980, 1984; Felsenstein 1981), additional sources of data can trigger effective, and informative, methods of inference for phenotypic traits (Pennell and Harmon 2013). Our results demonstrate how models consistent with a quantitative genetic interpretation can be identified statistically from other interpretations of OU models. Our comparisons of simulated *OU*_*Free*_ vs. *QG* models are somewhat contrived in that parameters were drawn directly using informative priors that were subsequently used as priors for model-fitting—an optimal scenario that is unrealistic when fitting real data. Furthermore, priors for the *QG* model were set to correspond to parameters estimated for body size data, a trait known to be unlikely to follow the *QG* model over million-year timescales (Lynch 1990; Hansen 1997; Butler and King 2004; Uyeda et al. 2011). Consequently, it is unsurprising we can obtain such dramatic support for the generating model when comparing *QG* and *OU*_*Free*_ parameterizations. Nevertheless, the *QG* model may be appropriate to apply to traits that are known to be more constrained, have lower additive genetic variances, if selection is known to be quite weak or if the underlying dynamics of adaptive landscapes are well-described by the model of peak movement. The potential for incorporating prior information is not limited to *QG* data. OU models have been effectively implemented to fit to fossil timeseries on timescales intermediate between microevolutionary, and phylogenetic scales (Hunt 2007, 2008, 2013; Reitan et al. 2012). The possibility of uniting these different data sources using either a single, phylogeny-based modeling framework, or by using the results of models fit to fossil data to inform the priors for comparative data (for example) provides rich avenue for unification of microevolutionary, fossil and macroevolutionary data.

Some may view the reversible-jump framework proposed in this study to be a data-mining tool that will lead to over-fitting of non-biologically relevant statistical noise. We agree that biologically informed *a priori* hypotheses should not be thrown out in any analysis and should be preferred to *a posteriori*, non-biologically based statistical models. However, we demonstrate one way to unite these approaches by comparing biologically informed *a priori* hypotheses to the posterior distribution of the unconstrained model. We tested the best-fitting model of a previous study (Jaffe et al. 2011) to demonstrate how *a priori* hypotheses can be compared to *a posteriori* hypotheses obtained in the reversible-jump framework. We find that the best-fitting model of chelonian carapace length evolution found in Jaffe et al. (2011) captures only one of the highly supported optima-shifts in the posterior distribution (Figure 6). The posterior obtained via reversible-jump inference is much more strongly supported based on Bayes Factors even given the high penalty assigned to models with vague priors (Kass 1993). Comparing biologically informed hypotheses to the posterior distribution of statistically supported hypotheses provides a useful metric for determining the adequacy of our model in explaining the data. Furthermore, we can generate additional hypotheses based on the inadequacy of the *a priori* hypotheses to explain the existence of distinct adaptive regimes for certain clades. For example, in the Chelonia dataset we examine here we find that there is a strongly supported adaptive shift in the softshell turtles (Trionychidae) that is not captured by the *OU*_*habitat*_ model of Jaffe et al. (2011). Based on this observation, we can conclude that the simple freshwater-marine dichotomy does not capture the underlying causal forces behind carapace length evolution. Instead, optimal body size may be better explained by other factors. For example, a shift to a larger phenotypic optimum may be accompanied by shifts to more aquatic lifestyles (irrespective of salinity) by releasing species from constraints imposed by the physical environment. Furthermore, it has been hypothesized that larger body sizes in chelonians may require higher environmental temperatures to enable high enough mass-specific metabolic rates to sustain growth (Makarieva et al. 2005; Head et al. 2009). Thus, a combination of an aquatic lifestyle and warm temperatures may favor shifts toward larger body sizes.

An important implication of our analysis is that the biological interpretation of a model may change the posterior distribution of model fits. For example, species of turtles may cluster broadly into adaptive zones defined based on aquatic or non-aquatic habits, and such transitions may be rare enough that OU model with a handful of shifts and weak parameters for *α* and *σ*^2^ could adequately describe evolutionary patterns. However, support for such a pattern does not exclude the possibility that at shorter timescales, species evolve according to a pulsed pattern of shifts in adaptive optima separate by million-year periods of phenotypic stasis [i.e. the pattern of stasis that lead to Eldredge and Gould’s (1972) proposal of punctuated equilibrium, but see Pennell et al. 2013]. If such a stasis model were enforced through informative priors, we would expect high values for *α* and *σ*, as well as more numerous shifts. Both evolution within these broad adaptive zones and intervals of stasis may be occurring simultaneously, and there may be statistical signals for both processes that are detectable in phylogenetic comparative data. 

~~~
bayou
~~~

 provides a flexible means of fitting these biological interpretations, and testing for specific processes and patterns of interest.

Our method helps alleviate many of the challenges inherent to fitting OU models to phylogenetic comparative data. Inference of OU models is often challenging due to ridges in likelihood space, which result in poor convergence and difficult to interpret parameter estimates. We emphasize the importance of examining the full posterior distribution, rather than point estimates, when interpreting model fits and the statistical signal of adaptive evolution in comparative data. For example, Ingram and Mahler’s 2013 

~~~
SURFACE
~~~

 method, as in bayou, searches for an optimal arrangement of adaptive regimes across the phylogeny. However, because of ridges in likelihood space in the chelonia dataset we examined, 

~~~
SURFACE
~~~

 gave highly unrealistic estimates of the values of the adaptive optima. This is because there are a range of correlated values of *α*, *σ*^2^ and *θ*_1_, …, *θ*_*K*_ that give essentially the same likelihood. Thus, the ML estimate in 

~~~
SURFACE
~~~

 combines extremely distant optima (ranging from −422.6 to 111.2; Table 3) with much weaker estimates of *α* (phylogenetic half-life of 92.6 vs. 3.94 in bayou) to obtain very high log-likelihoods relative to other methods (Table 3, Figure 6). bayou avoids these difficult to interpret results by using informative prior information to exclude biologically unreasonable models. Although this introduces some subjectivity, most biologists would agree that we can reasonably reject the idea that any extant species of chelonians are adapting to an optimal carapace length of *e*^111.2^ = 1.96*x*10^43^ km long (i.e. larger than the diameter of the observable universe). Since fitted model parameters are correlated, unreasonable estimates of adaptive optima also affect estimation of *α*, *σ*^2^ and the location of shifts. By simply setting a reasonable prior on this one set of parameters (*θ*_1_, …, *θ*_*K*_), we show that the estimation of *α* and *σ* collapses from a ridge to a narrow peak of values (Figure 7). Thus, while both methods can infer the location of shifts in adaptive optima without *a priori* specification, bayou allows for the inclusion of informative priors and returns a full posterior of credible models; while 

~~~
SURFACE
~~~

 allows for convergent regimes (the “backwards” selection step) which is currently not implemented in the reversible-jump framework of 

~~~
bayou
~~~

.

Our analysis of simulations results gives considerable insight into best practices in fitting OU models to phylogenetic comparative data. While 

~~~
bayou
~~~

 does not estimate the number of shifts with much power, it is nonetheless possible to distinguish models with little statistical support for adaptive peak shifts vs. models with strong evidence of optima shifts. For example, as the true model approaches BM (i.e. high phylogenetic half-lives) or the number of shifts goes to 0, the posterior support for any particular branch greatly decreases, eventually reaching the probability of a random branch having a shift given the prior density (Figure 1; Online Appendix III). Thus, if no branch has high posterior support relative to the prior density, we conclude there is little evidence for an adaptive shift (regardless of the mean number of shifts inferred in the posterior distribution). In fact, there is relatively little cost to placing a high prior on the number of shifts for inference of phylogenetic half-life, *σ*^2^, and the location and magnitude of optima shifts, whereas conservative priors will often result in mis-estimation of these parameters if the true model is complex. Therefore, we recommend using priors that favor a moderate to large number of shifts rather than using more conservative priors (see Online Appendix III). Furthermore, if the posterior for phylogenetic half-life includes values higher than the phylogenetic tree height, this indicates that the model is very BM-like and it is unlikely that shifts will be estimated. This is because optima shifts are expected to only weakly affect the distribution of the data. As a simple heuristic, we obtained reasonable estimates of parameters when the phylogenetic half-life is between the youngest split in the phylogeny and the total tree height. Half-life values substantially less than the youngest split in the phylogeny indicate that after accounting for optima shifts, residual variation within regimes is not phylogenetically correlated (white noise). By contrast, phylogenetic half-lives much higher than tree height indicate strong phylogenetic signal even after accounting for shared adaptive optima, which is suggestive of BM-like evolution.

A number of extensions are possible within our proposed method, and is a first step toward a much more expansive suite of models. Using likelihood approaches, OU models have been expanded to include randomly evolving continuous predictors (Hansen et al. 2008), multivariate evolution of correlated traits (Bartoszek et al. 2012), varying *α* and *σ*^2^ parameters across the tree (Beaulieu et al. 2012; Lapiedra et al. 2013) and identification of convergent regimes (Ingram and Mahler 2013; Mahler et al. 2013). Expanding these models to a Bayesian framework would carry many of the same advantages we describe in this study. Furthermore, we emphasize the importance of developing models that describe the dynamics of adaptive landscapes themselves, and suggest anchoring these models in empirical datasets through the use of informative priors will greatly improve our understanding of macroevolutionary dynamics.

## Acknowledgements

We thank Jon Eastman for useful discussion and for providing code upon which 

~~~
bayou
~~~

 was built. We would like to thank Matt Pennell, Paul Joyce, and the Harmon lab for useful discussions. Tanja Stadler and two anonymous reviewers provided useful comments on an earlier version of this manuscript. This work was supported by NSF (DEB 091499, DEB 1208912 and DBI 0939454).

